## Supplementary material for "Presynaptic Rac1 controls synaptic strength through the regulation of synaptic vesicle priming": Fig6 - Figure supplement 1

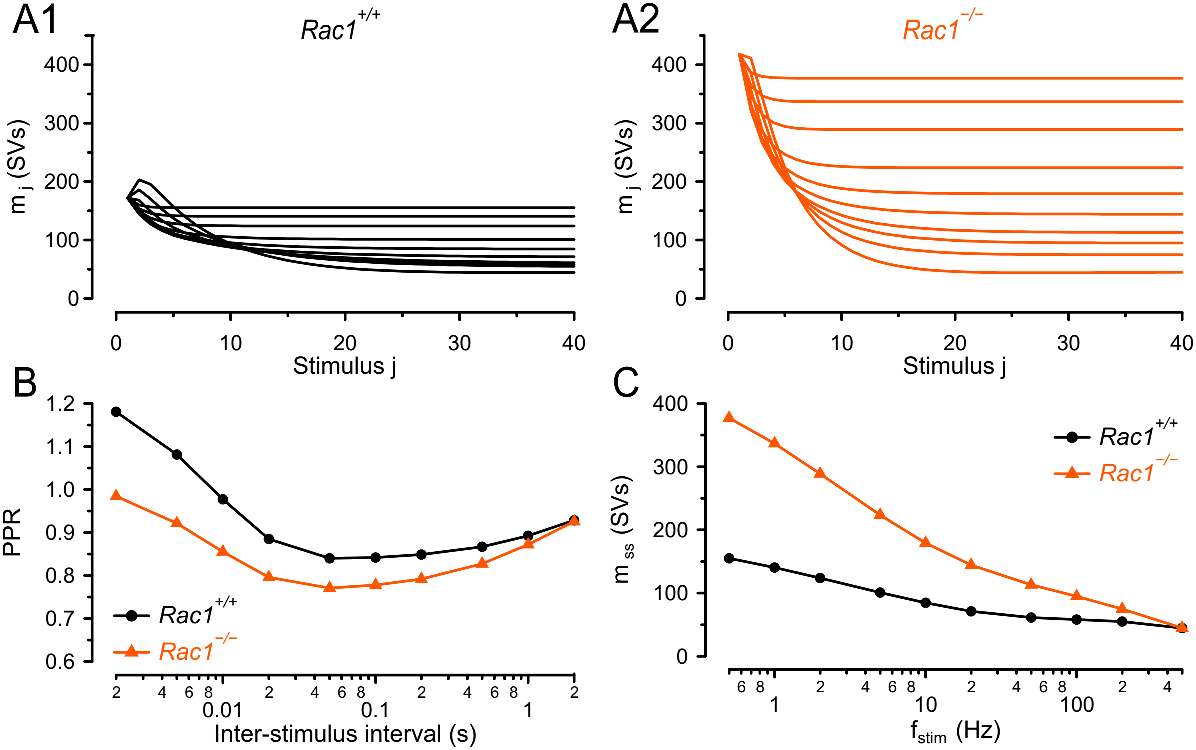


D1

| Parameter | Value | Unit |
|---|---|---|
| $[Ca^{2+}]$ decay τ | 0.028 | s |
| $[Ca^{2+}]$ amplitude | 1.0E-07 | M |
| $P_r$ | 0.08 | |
| $N_{total}$ | 2650 | |
| $k_f$ | 0.5 | $s^{-1}$ |
| $k_b$ | 0.116 | $s^{-1}$ |
| σ | 9.6E+06 | $s^{-1}\ M^{-1}$ |
| $K_{0.5}$ | 3.1E-06 | M |
| $y_{inc}$ | 0.5 | |
| $z_{dec}$ | 0.4 | |
| $y_{max}$ | 1.2 | |
| $z_{min}$ | 0.75 | |
| $\tau_y$ | 0.01 | s |
| $\tau_z$ | 3 | s |

D2

| Parameter | Value | Unit |
|---|---|---|
| $[Ca^{2+}]$ decay τ | 0.028 | s |
| $[Ca^{2+}]$ amplitude | 1.0E-07 | M |
| **$P_r$** | **0.165** | |
| **$N_{total}$** | **2900** | |
| **$k_f$** | **0.8** | **$s^{-1}$** |
| $k_b$ | 0.116 | $s^{-1}$ |
| **σ** | **1.8E+07** | **$s^{-1}\ M^{-1}$** |
| **$K_{0.5}$** | **6.6E-07** | **M** |
| $y_{inc}$ | 0.5 | |
| $z_{dec}$ | 0.4 | |
| **$y_{max}$** | **1.15** | |
| $z_{min}$ | 0.75 | |
| $\tau_y$ | 0.01 | s |
| $\tau_z$ | 3 | s |
