## Supplementary material for "Presynaptic Rac1 controls synaptic strength through the regulation of synaptic vesicle priming": Fig6 - Figure supplement 2

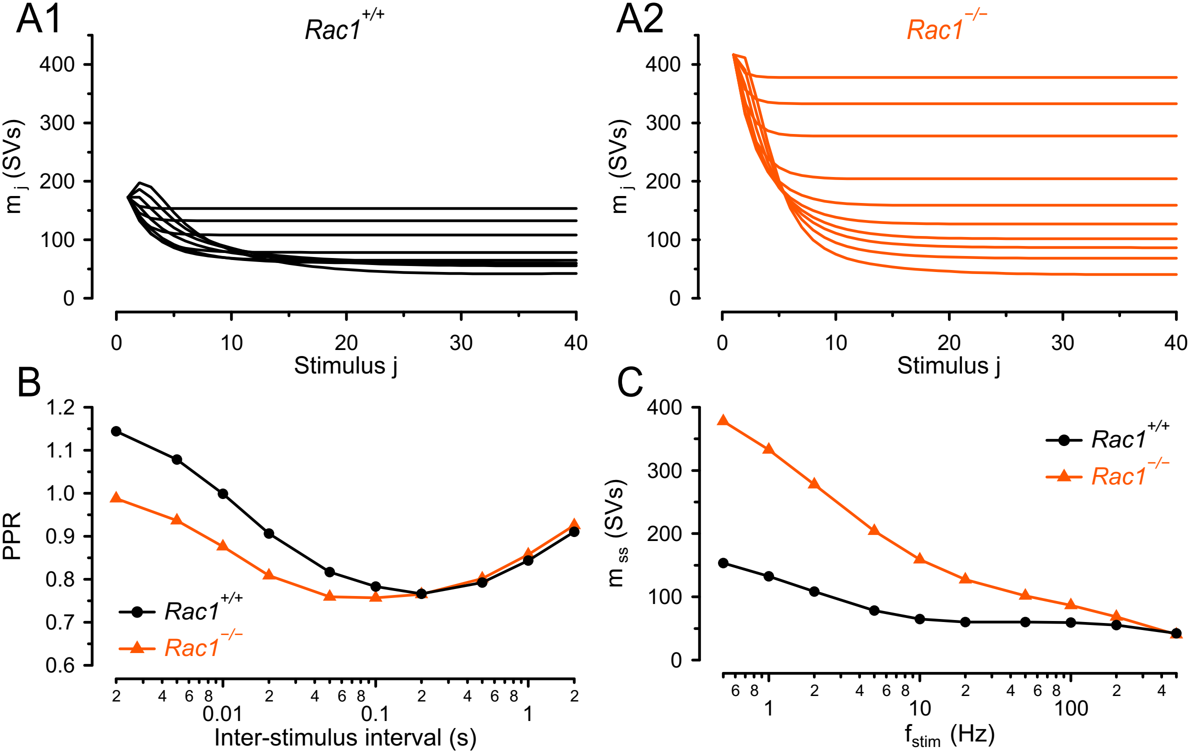


D1

| Parameter | Value | Unit |
|---|---|---|
| $[Ca^{2+}]$ decay τ | 0.043 | s |
| $[Ca^{2+}]$ amplitude | 9.5E-08 | M |
| $P_r$ | 0.25 | |
| $N_{total}$ | 3000 | |
| $k_1$ | 0.84 | $s^{-1}$ |
| $b_1$ | 0.42 | $s^{-1}$ |
| $\sigma_1$ | 5.7E+06 | $s^{-1}\,M^{-1}$ |
| $k_2$ | 0.31 | $s^{-1}$ |
| $b_2$ | 0.69 | $s^{-1}$ |
| $\sigma_2$ | 4.6E+06 | $s^{-1}\,M^{-1}$ |
| TSL decay τ | 0.08 | |
| TSL fraction | 0.08 | |
| $K_{0.5}$ | 5.6E-06 | M |
| $y_{inc}$ | 0.34 | |
| $z_{dec}$ | 0.4 | |
| $y_{max}$ | 1.23 | |
| $z_{min}$ | 0.8 | |
| $\tau_y$ | 0.014 | |
| $\tau_z$ | 3 | |

D2

| Parameter | Value | Unit |
|---|---|---|
| $[Ca^{2+}]$ decay τ | 0.043 | s |
| $[Ca^{2+}]$ amplitude | 9.5E-08 | M |
| $P_r$ | 0.25 | |
| $N_{total}$ | 3000 | |
| **$k_1$** | **1.05** | **$s^{-1}$** |
| **$b_1$** | **0.35** | **$s^{-1}$** |
| **$\sigma_1$** | **1.2E+07** | **$s^{-1}\,M^{-1}$** |
| **$k_2$** | **1.3** | **$s^{-1}$** |
| **$b_2$** | **0.78** | **$s^{-1}$** |
| **$\sigma_2$** | **2.2E+07** | **$s^{-1}\,M^{-1}$** |
| TSL decay τ | 0.08 | |
| TSL fraction | 0.08 | |
| **$K_{0.5}$** | **8.5E-07** | **M** |
| $y_{inc}$ | 0.34 | |
| $z_{dec}$ | 0.4 | |
| $y_{max}$ | 1.23 | |
| $z_{min}$ | 0.8 | |
| $\tau_y$ | 0.014 | |
| $\tau_z$ | 3 | |
